# *Leishmania major* degrades murine CXCL1 – an immune evasion strategy

**DOI:** 10.1101/571141

**Authors:** Matthew S. Yorek, Barun Poudel, Lalita Mazgaeen, R. Marshall Pope, Mary E. Wilson, Prajwal Gurung

## Abstract

Leishmaniasis is a global health problem with an estimated report of 2 million new cases every year and more than 1 billion people at risk of contracting this disease in endemic areas. The innate immune system plays a central role in controlling *L. major* infection by initiating a signaling cascade that results in production of pro-inflammatory cytokines and recruitment of both innate and adaptive immune cells. Upon infection with *L. major*, CXCL1 is produced locally and plays an important role in the recruitment of neutrophils to the site of infection. Herein, we report that *L. major* specifically targets murine CXCL1 for degradation. The degradation of CXCL1 is not dependent on host factors as *L. major* can directly degrade recombinant CXCL1 in a cell-free system. Using mass spectrometry, we discovered that the *L. major* protease cleaves at the C-terminal end of murine CXCL1. Finally, our data suggest that *L. major* metalloproteases are involved in the direct cleavage and degradation of CXCL1, and a synthetic peptide spanning the CXCL1 cleavage site can be used to inhibit *L. major* metalloprotease activity. In conclusion, our study has identified an immune evasion strategy employed by *L. major* to evade innate immune responses in mice, likely reservoirs in the endemic areas, and further highlights that targeting these *L. major* metalloproteases may be important in controlling infection within the reservoir population and transmittance of the disease.

**Authors’ summary:** Our study discovered a highly specific role for *L. major* metalloprotease in cleaving and degrading murine CXCL1. Indeed, *L. major* metalloprotease did not cleave murine CXCL2 or human CXCL1, CXCL2 and CXCL8. CXCL1 is a critical chemokine required for neutrophil recruitment to the site of infection; thus, we propose that this metalloprotease may have evolved to evade immune responses specifically in the murine host. We have further identified that the C-terminal end on CXCL1 is targeted for cleavage by the *L. major* metalloprotease. Finally, this cleavage site information was used to design peptides that are able to inhibit CXCL1 degradation by *L. major*. Our study highlights an immune evasion strategy utilized by *L. major* to establish infection within a murine host.

## Introduction

*Leishmania spp*. are unicellular eukaryotic protozoan parasites that are transmitted to mammalian hosts by sandfly (*Phlebotomine and Lutzomyia spp*.) bites [1]. Upon transmission of *L. major* promastigotes (the infectious stage for mammalian hosts with a long slender body and an anterior flagellin), the promastigotes are quickly taken up by neutrophils, macrophages and keratinocytes [2–6]. Within the macrophages, *Leishmania spp*. promastigotes hijack the phagocytic vacuole and transform into amastigotes (round body lacking an anterior flagellin) [7]. The *Leishmania spp*. amastigotes then proliferate within the vacuole and establish infection within the host [8, 9]. While a mammalian host-vector system is the major mode of *Leishmania spp*. transmission, several studies have reported a vertical transmission of these parasites in mammalian hosts, from a pregnant female to its offspring [10–13]. Specifically, *Leishmania spp*. infection has been found to be endemic in foxhound dog populations in the United States, where the vectors are not present [14–17]. Given that *Leishmania spp*. can infect several hosts, including rodents and dogs (in addition to humans) [18], these studies demonstrate how these parasites can remain endemic even with strategies to eradicate sandflies.

Our immune system is extremely efficient in killing pathogens. Professional phagocytes such as macrophages and neutrophils phagocytose and kill the invading pathogen in the intracellular phagosome-lysosome compartment. In addition, these phagocytes also respond to the foreign pathogens by secreting pro-inflammatory cytokines and chemokines to recruit neutrophils to the site of infection [19–21]. *Leishmania spp*. have evolved to evade the host immune response by using its armada of virulence factors to avoid host killing [22]. The two major virulence factors of *Leishmania spp*. include leishmanolysin metalloprotease glycoprotein 63 (GP63) and lipophosphoglycan (LPG), and both have been extensively studied for their roles in immune evasion [23, 24]. GP63 and LPG inhibit the formation of the membrane attack complex to evade complement-mediated lysis [25, 26], inhibit acidification of leishmania containing vacuoles [27, 28], and dampen host immune signaling pathways [29, 30] for establishing infection within the mammalian host. In addition, *Leishmania spp*. can hijack the host immune responses to establish infections as shown by *Leishmania chagasi (L. chagasi)*-mediated activation of TGF-β and *Leishmania major (L. major)*-induced activation of NLRP3 inflammasome, events that promote *L. major* survival and pathology [31–33]. Here, we have identified one such immune evasion mechanism employed by *L. major*.

*L. major* is the most common *Leishmania spp*. and a major cause of cutaneous leishmaniasis that affects 600,000-1,000,000 people globally [34]. Once deposited at the site of a sandfly bite, *L. major* promastigotes are taken up by the resident macrophages and keratinocytes, which then secrete essential cytokines and chemokines to elicit an innate immune response [35]. One of the earliest chemokines that are produced in the skin in response to *L. major* include CXCL1 [36]. CXCL1 is a functional homolog of human IL-8 in mice and a potent neutrophil chemoattractant [37–41]. Keratinocytes [3], neutrophils [42] and macrophages [43, 44] have all been suggested to produce CXCL1 (or IL-8 in humans) in response to *Leishmania spp*. infection. Several studies have shown that neutrophils infiltrate the site of *Leishmania spp*. infection as early as 1-hour post-infection and are important for optimal resolution of the infection [21, 45]. Given the importance of CXCL1 in modulating the early innate immune response, we reasoned that *L. major* targets CXCL1 to evade early host innate immune responses.

In this current study, we have identified murine CXCL1 as a highly specific substrate for *L. major* metalloprotease and a possible immune evasion strategy employed by this parasite to establish a successful infection within the murine host. We further report that *L. major* promotes proteolytic cleavage of murine CXCL1 at the C-terminal end to initiate its degradation. Finally, we have designed a synthetic peptide spanning the CXCL1 cleavage site that inhibits *L. major* protease activity. In conclusion, our study has uncovered a specific mechanism employed by *L. major* to degrade murine CXCL1 which may help the parasite to establish infection within the murine host.

## Results

### *L. major* infection abrogates LPS-induced CXCL1 production by bone marrow-derived macrophages

To investigate how *L. major* impacts innate immune responses elicited by macrophages, we stimulated bone marrow-derived macrophages (BMDM) with lipopolysaccharide (LPS) in the presence or absence of *L. major* (WHOM/IR/-173) infection following established protocol [32]. As detailed in the experimental outline (Fig. 1A), BMDM were stimulated with 20 ng/ml LPS in the presence or absence of 20 MOI *L. major* promastigotes for 48 hours. LPS stimulation of BMDM induced robust production of IL-6, TNF and CXCL1. Simultaneous LPS stimulation and *L. major* infection did not impact the production of IL-6 and TNF by BMDM whereas CXCL1 levels were significantly blunted (Fig. 1A). These data suggest that *L. major* specifically targets LPS-induced CXCL1 production by BMDM.

**Figure 1:**
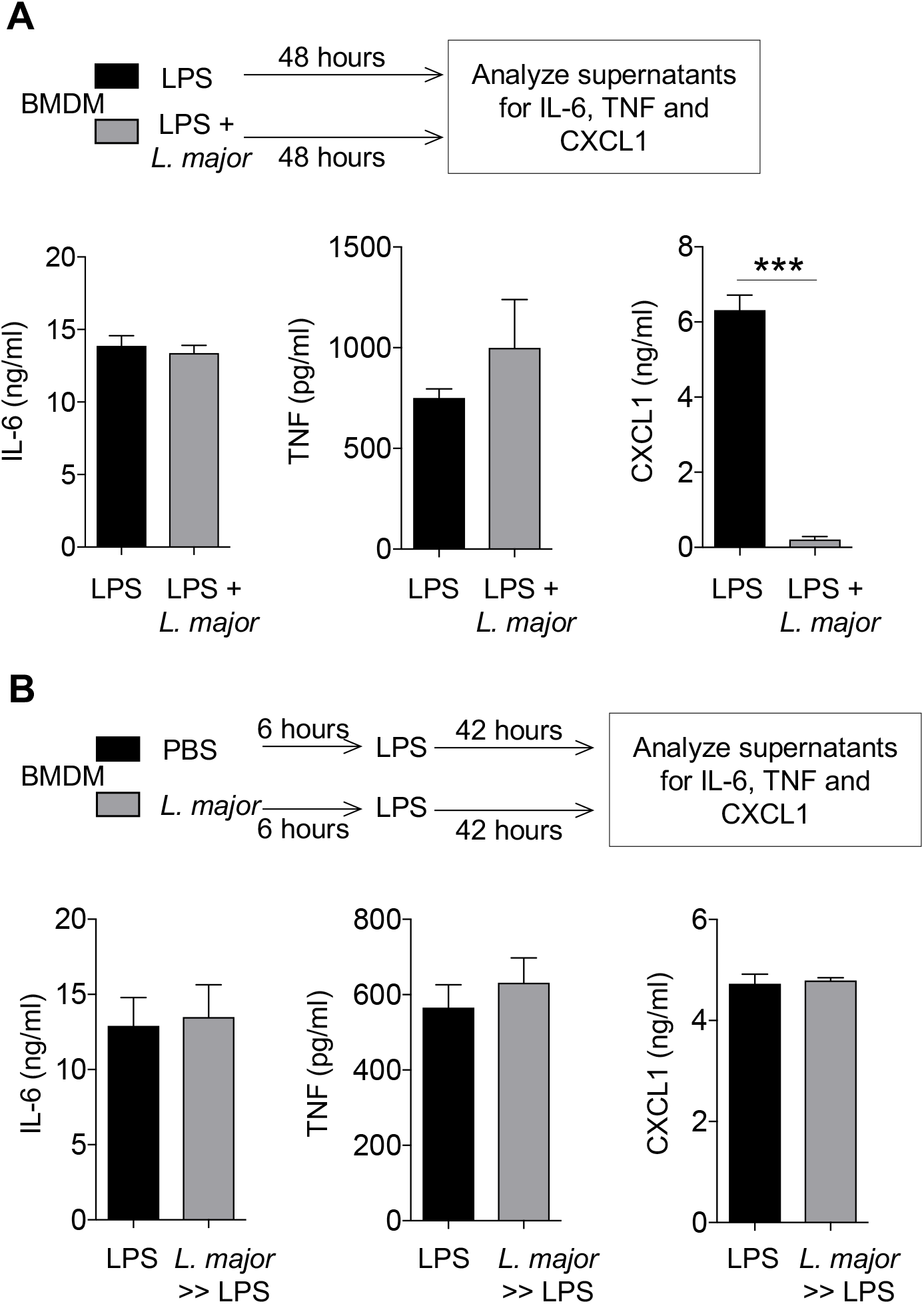
*L. major* diminishes LPS-induced CXCL1 release by BMDMs. (A) BMDMs were treated with LPS (20 ng/ml) in the presence or absence of *L. major* (20 MOI) for 48h and cell culture supernatants were analyzed for the indicated cytokines. (B) BMDMs were treated with PBS or *L. major* (20 MOI) for 6h. Then, the cells were washed to remove extracellular *L. major* and stimulated with LPS for next 42h and cell culture supernatants were subjected to cytokines analysis as in (A). Data are representative of at least three independent experiments. Results are represented as mean ± SEM. \*\*\**P*<0.001.

Previous studies have shown that *L. major* targets signaling pathways to evade immune responses and establish infection [[29]]. To examine whether *L. major* specifically targeted CXCL1 expression and production, we designed an experiment whereby BMDM were infected with 20 MOI *L. major* for 6 hours, extracellular *L. major* were washed, then stimulated with 20 ng/ml LPS for the next 42 hours (Fig. 1B). Interestingly, *L. major*-infected BMDM produced equal levels of CXCL1 when compared to controls (Fig. 1B). These results show that inhibition of CXCL1 by *L. major* may not occur through modulation of intracellular signaling pathway that promotes CXCL1 expression and/or production by BMDM.

### CXCL1 detection is significantly reduced by *L. major* in a cell free system

Given that *L. major* pre-infection of BMDM prior to LPS stimulation did not affect CXCL1 production, we posited that *L. major* regulates secreted CXCL1 in the extracellular milieu. To this end, we used supernatants derived from LPS-stimulated BMDM as a source of CXCL1 and cultured 500 μl of these cell-free supernatants with 20 × 10^6^ *L. major* promastigotes for 24 hours (Fig. 2A). While levels of IL-6 and TNF detected in the supernatants remained unchanged, levels of CXCL1 were significantly reduced after addition of *L. major* to the supernatants. The reduced levels of CXCL1 were not due to its short half-life in the culture, as the levels of CXCL1 in the cell free supernatants were stable up to 48 hours (Supplemental Fig. 1). Thus, *L. major* directly dampens CXCL1 detection.

**Figure 2:**
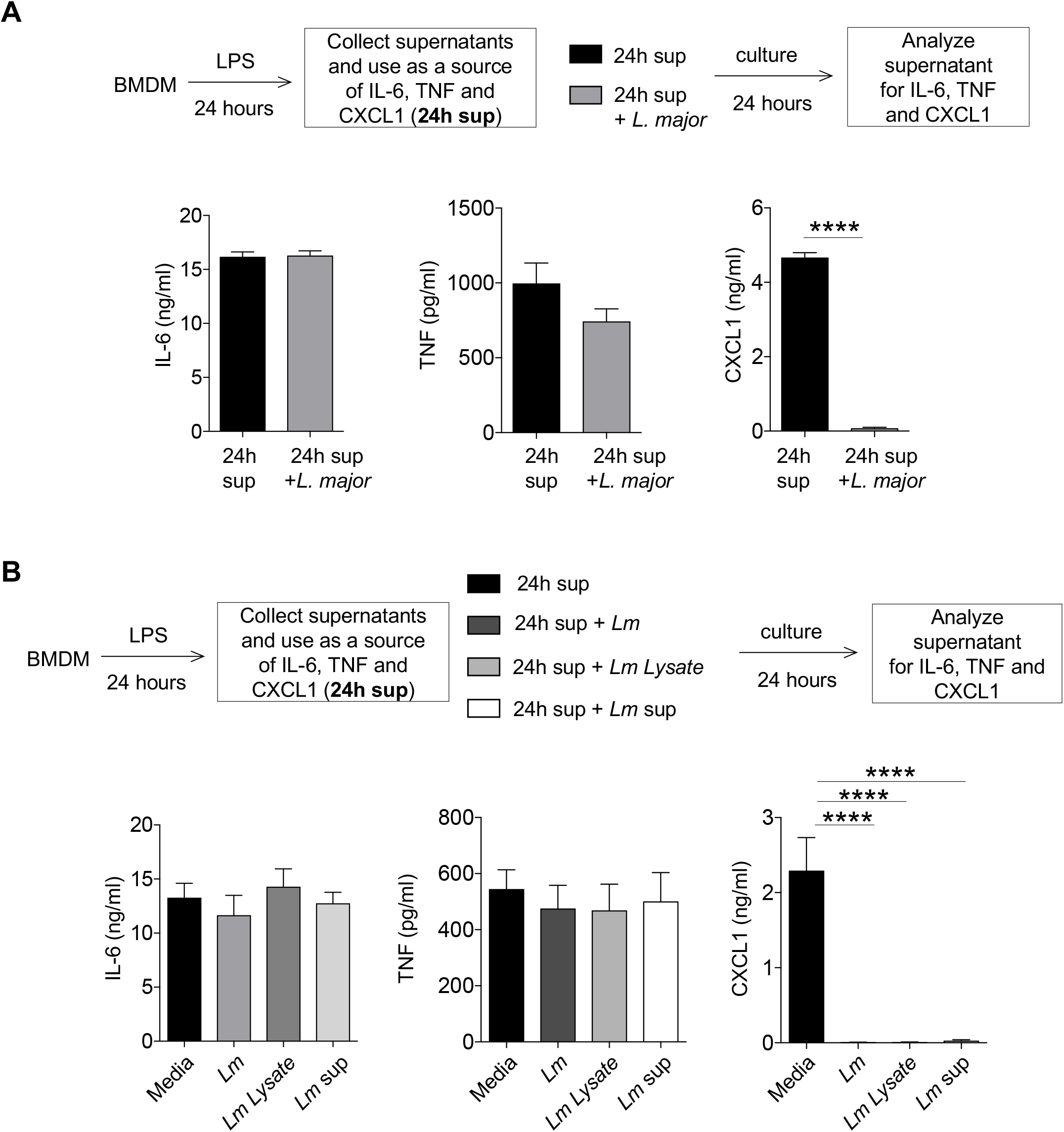
*L. major* inhibits CXCL1 detection in a cell free system. (A) Conditioned supernatants from 24h LPS (20 ng/ml)-stimulated BMDMs were treated with or without *L. major* for 24h and cytokines were analyzed by ELISA. (B) 24h LPS treated conditioned media from BMDMs were treated with or without *L. major, L. major* lysate (*Lm* lysate), or *L. major* culture supernatant (*Lm* sup) for 24h and cytokines were measured as in (A). Data are representative of at least three independent experiments. Results are represented as mean ± SEM. \*\*\*\**P*<0.0001.

We next examined whether the observed effect of *L. major* on CXCL1 detection was dependent on live parasites. To this end, we cultured supernatants from LPS-stimulated BMDM with *L. major* lysates (*Lm* lysate: generated by 3× freeze-thaw cycles of *L. major*) or supernatants from *L. major* culture (*Lm* sup: supernatant collected from stationary phase of *L. major* growth; Fig. 2B). Similar to the addition of live *L. major*, addition of *Lm* lysate or *Lm* sup also reduced CXCL1 in the supernatants while IL-6 and TNF remained unaffected (Fig. 2B). These results demonstrate that: first, *L. major* need not be alive to dampen CXCL1 detection; second, *L. major* lysate can dampen CXCL1 detection; and finally, *L. major* secreted factors dampen CXCL1 detection. Boiling *Lm* lysate or *Lm* sup for 20 minutes at 100°C rescued CXCL1 detected in the supernatants, suggesting that the responsible *L. major* components are susceptible to heat treatment (Supplemental Fig. 2). Given the sensitivity to heat treatment, we propose that the CXCL1 regulating *L. major* components are proteinaceous.

### *L. major* reduces recombinant murine CXCL1 (rm-CXCL1) levels in a cell free system

Macrophages secrete several hundreds of different proteins that may indirectly alter our observed effect of *L. major* components on CXCL1 [46]. To this end, we obtained recombinant murine CXCL1 (rm-CXCL1, Tonbo Biosciences, San Diego, CA) which was stable in culture up to 48 hours (Supplemental Fig. 3). Importantly, *Lm* sup addition to rm-CXCL1 dampened its detection by ELISA in a time-dependent manner, and boiling *Lm* sup rescued rm-CXCL1 detection (Supplemental Fig. 3).

To further investigate the specificity of *L. major* in reducing rm-CXCL1 levels, we examined rm-CXCL2, rh-CXCL1, rh-CXCL2 and rh-CXCL8 sequence homology by CLUSTAL-W alignment and found significant homology between these recombinant proteins. Despite the significant homology, *L. major* failed to reduce levels of rm-CXCL2, rh-CXCL1, rh-CXCL2 or rh-CXCL8, demonstrating specific regulation of rm-CXCL1 (Fig. 3, B-F). As expected, *L. major* did not inhibit rm-TNF levels (Fig. 3G). Given the specific nature of *L. major* (WHOM/IR/-170) in dampening rm-CXCL1 levels, we next determined whether this activity was specific to the WHOM/IR/-170 isolate. However, supernatant from *L. major* (IA0, isolated from a patient in Iowa who acquired *L. major* in Iraq [47]) was also able to inhibit rm-CXCL1 detection when compared to the WHOM/IR/-170 strain (Fig. 3H). Furthermore, exosomes from the *L. major* supernatants from both isolates were able to reduce rm-CXCL1 levels, suggesting that the active components are also present in the *L. major*-derived vesicles (Fig. 3H).

**Figure 3:**
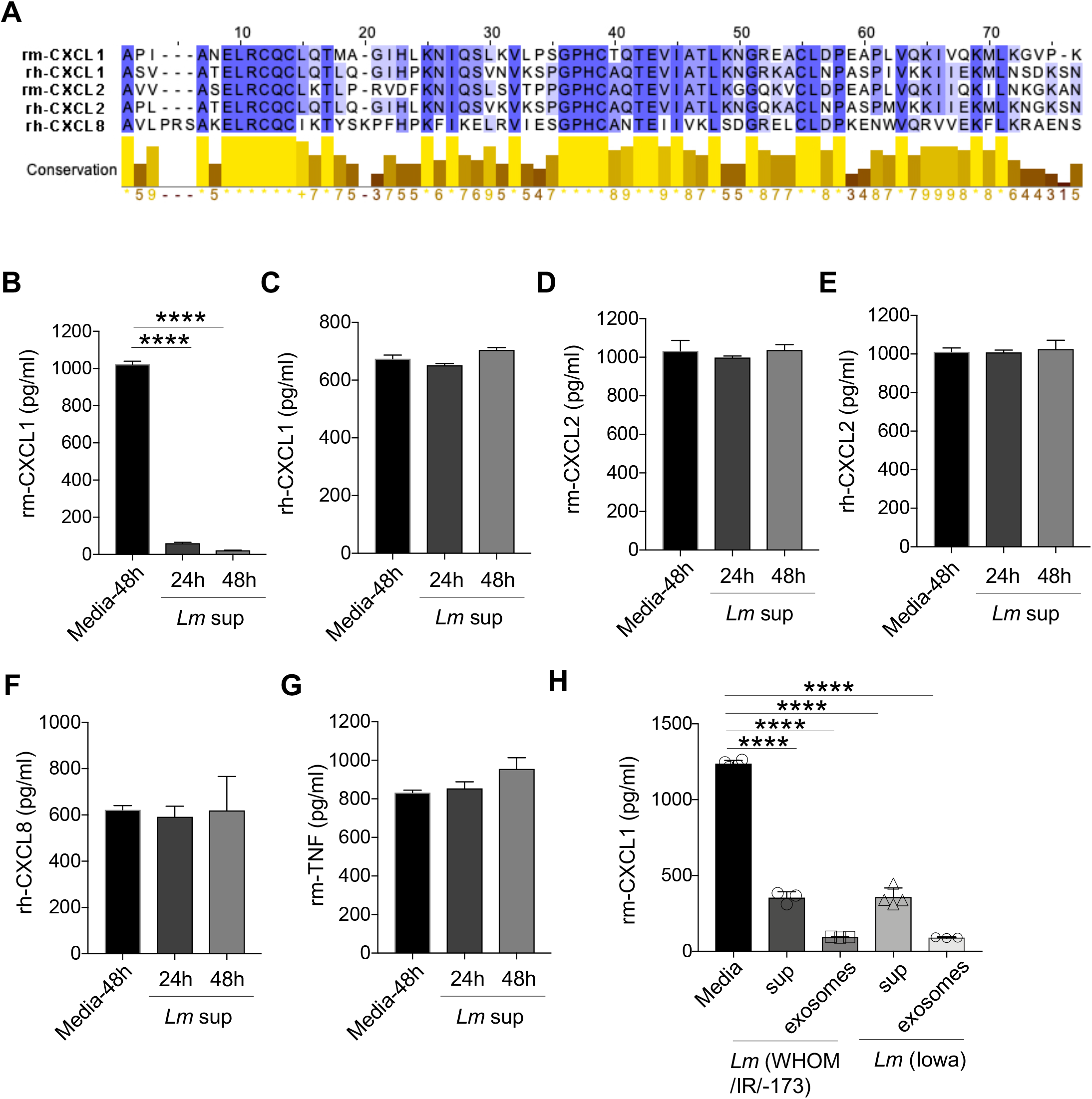
Selective inhibitory activity of *L. major* on recombinant murine CXCL1. (A) Sequence alignment of recombinant murine (rm) and recombinant human (rh) CXCL1, CXCL2, and CXCL8. (B) Rm-CXCL1, (C) rh-CXCL1, (D) rm-CXCL2, (E) rh-CXCL2, (F) rh-CXCL8, and (G) rm-TNF were incubated with or without *Lm* sup for indicated time-points and ELISA was performed to quantify the mentioned cytokine/chemokines. (H) Rm-CXCL1 was incubated with or without *Lm* sup or *Lm* exosomes derived from indicated strains of *L. major* for 24h, and CXCL1 quantification was performed as in (G). Data are representative of at least three independent experiments. Results are represented as mean ± SEM. \*\*\*\**P*<0.0001.

### *L. major* inactivates biological activity of CXCL1

Our data show that *L. major* reduced CXCL1 levels as demonstrated by ELISA (Figs. 1–3). Possible reasons why CXCL1 levels were reduced when incubated with *L. major* include: 1) *L. major* proteins bind to CXCL1, limiting the ability of anti-CXCL1 antibody in ELISA to interact with CXCL1 or 2) *L. major* proteases cleaves and degrades CXCL1. Considering both outcomes, it is possible that CXCL1, either masked or cleaved by *L. major* components, could still be biologically active. To this end, we performed a functional assay where rm-CXCL1 or rm-CXCL1+Lm lysate was used to stimulate BMDM (Fig. 4A). As demonstrated previously, when rm-CXCL1 is incubated with *L. major*, rm-CXCL1 levels are significantly reduced (Fig. 4B). *Lm* lysate alone has very little stimulatory activity as demonstrated by mRNA induction of *Cxcl1, Tnf* and *Il6* (Fig. 4, C-E). As expected, rm-CXCL1 induced modest increase of *Cxcl1, Tnf* and *Il6* but rm-CXCL1+Lm lysate failed to induce *Cxcl1, Tnf* and *Il6* mRNA (Fig. 4, C-E). These results altogether demonstrate that *L. major* inactivates biological activity of rm-CXCL1.

**Figure 4:**
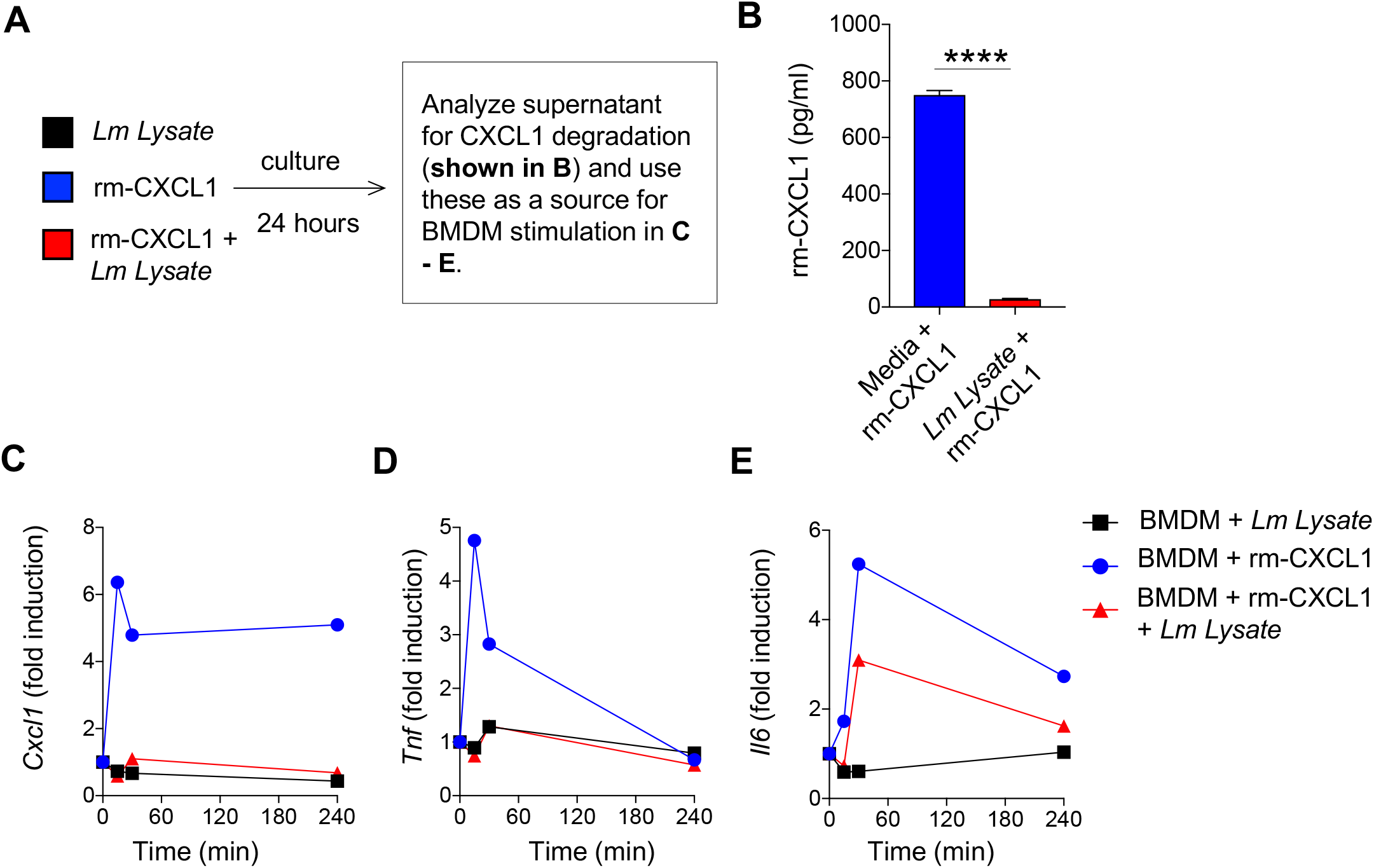
Biological activity of *L. major*-treated CXCL1 is diminished. (A) Schematic representation of experimental design. (B) Recombinant murine CXCL1 was incubated with *Lm* lysate for 24h and CXCL1 levels was determined by ELISA. (C) BMDMs were treated *Lm* lysate, rm-CXCL1, or *Lm* lysate + rm-CXCL1 from (B) for indicated time-points and mRNA expression of pro-inflammatory genes were evaluated by qPCR. Data are representative of at least three independent experiments.

### *L. major* cleaves CXCL1 at the C-terminal end to promote its degradation

To determine whether CXCL1 is degraded by *L. major*, we incubated rm-CXCL1 with *Lm* lysate for various time periods and examined rm-CXCL1 by silver staining (Fig. 5A). Rm-CXCL1 was detected at approximately 10KDa by silver stain. Interestingly, when incubated with *Lm* lysate rm-CXCL1 showed two bands as early as 30 minutes after incubation, suggesting *Lm* lysate-mediated cleavage of rm-CXCL1 (Fig. 5A and Supplemental Fig. 4). By 2 hours, only the cleaved rm-CXCL1 was observed and this cleaved band intensity decreased over time, suggesting further degradation (Fig. 5A). *L. major*-mediated cleavage of rm-CXCL1 was specific because similar experiments done with rm-CXCL2, rh-CXCL1 and rh-CXCL2 did not result in cleavage or degradation of these recombinant proteins (Fig. 5B).

**Figure 5:**
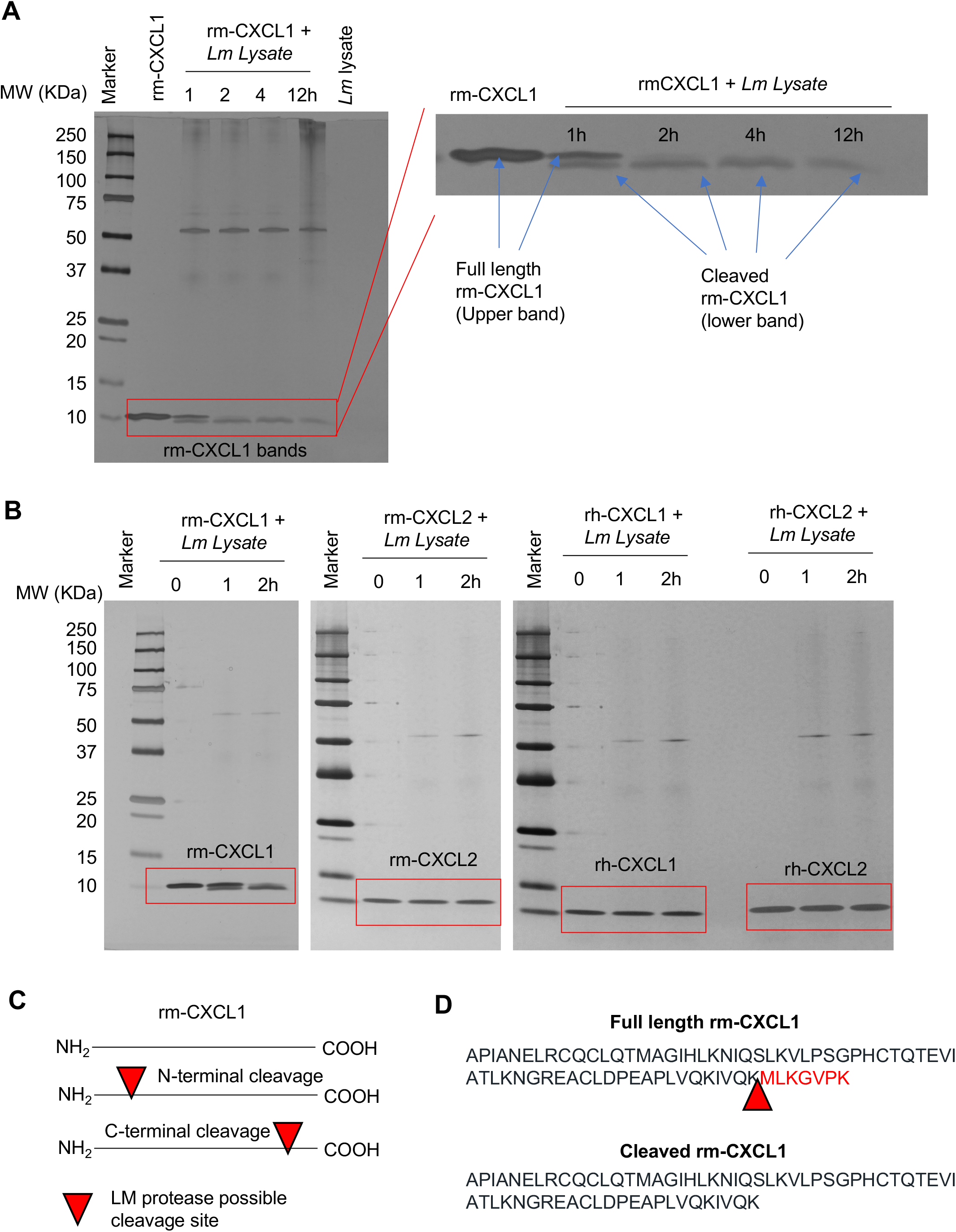
*L. major* cleaves CXCL1 at the C-terminal end. (A) A silver-stained SDS-PAGE demonstrate time-dependent cleavage of rm-CXCL1. Rm-CXCL1 were left alone or treated with *Lm* lysate for the indicated time points. Rm-CXCL1 bands were enlarged to show the cleavage products on right. (B) Indicated chemokines were treated with *Lm* lysate for indicated time-points and subjected to SDS-PAGE followed by silver-staining. (C) Hypothesized *L. major* cleavage site on rm-CXCL1 based on the observed cleaved rm-CXCL1 bands in (A and B). Arrow denotes possible cleavage site on CXCL1 (D) Identification of CXCL1 cleavage site by Mass Spectrometry analysis (raw data presented in Supplemental Fig. 4). Arrow represents cleavage site on rm-CXCL1 targeted by *L. major*. Data in A and B are representative of at least three independent experiments.

Based on the cleavage pattern of rm-CXCL1 (less than 1KDa shift), cleavage either at the N-terminal or C-terminal end would result in a large fragment (detected by silver stain as the lower band, Fig. 5A) and a smaller fragment (which could not be detected by silver stain; Fig. 5C). To identify the cleavage product and site, we processed the full-length rm-CXCL1 and *Lm* lysate-cleaved rm-CXCL1 bands by trypsin digestion and performed mass spectrometry to determine the sequence of the bands (Supplemental Fig. 5). The tryptic peptide sequence readouts of full-length rm-CXCL1 covered the whole rm-CXCL1 peptide sequence (Supplemental Fig. 5B), while the peptide coverage of cleaved rm-CXCL1 covered all of the rm-CXCL1 peptide sequence except the last 7 amino acids (MLKGVPK) at the C-terminal end (Supplemental Fig. 5C). Thus, *L. major* cleaves rm-CXCL1 after lysine 65 (K65) residue that results in a large N-terminal rm-CXCL1 fragment lacking 7 amino acids at the C-terminal end (Fig. 5D).

### *L. major* metalloprotease cleaves CXCL1

*L. major* has several proteases that enable it to survive within a cell and establish infection [48]. Metalloproteases and cysteine, serine and aspartic proteases are the major proteases described in *L. major* [48]. To examine whether cleavage activity was dependent on proteases, we first treated our rm-CXCL1 + *Lm* sup culture with pan protease inhibitor (Roche and Sigma) (Fig.6, A-C). Roche cOmplete protease inhibitor is a broad inhibitor of proteases including serine, cysteine and metalloproteases, and Sigma P8340 inhibitor is reported to inhibit serine, cysteine, acid proteases and aminopeptidases. However, the presence of protease inhibitors from Roche (cOmplete) or Sigma (P8340) did not inhibit *Lm* sup-mediated degradation of rm-CXCL1 as demonstrated by ELISA (Fig. 6, B and C, and Supplemental Fig. 6). Marimastat, a broad inhibitor of matrix metalloproteases [49], did not rescue the degradation of rm-CXCL1 by *Lm* sup (Supplemental Fig. 6B). EDTA treatment, which chelates metal ions such as Ca^2+^ and Fe^3+^ (and thus can inhibit certain metalloprotease), did not rescue rm-CXCL1 cleavage by *Lm* sup (Fig. 6, D and E). Interestingly, 1,10-Phenanthroline, a Zn^2^+ metalloprotease inhibitor [50], rescued rm-CXCL1 degradation by *Lm* sup (Fig. 6, F and G). While the addition of 1,10-Phenanthroline resulted in the upward shift of rm-CXCL1 bands in SDS-PAGE gels; importantly, no cleavage or degradation of rm-CXCL1 was observed (Fig. 6G).

**Figure 6:**
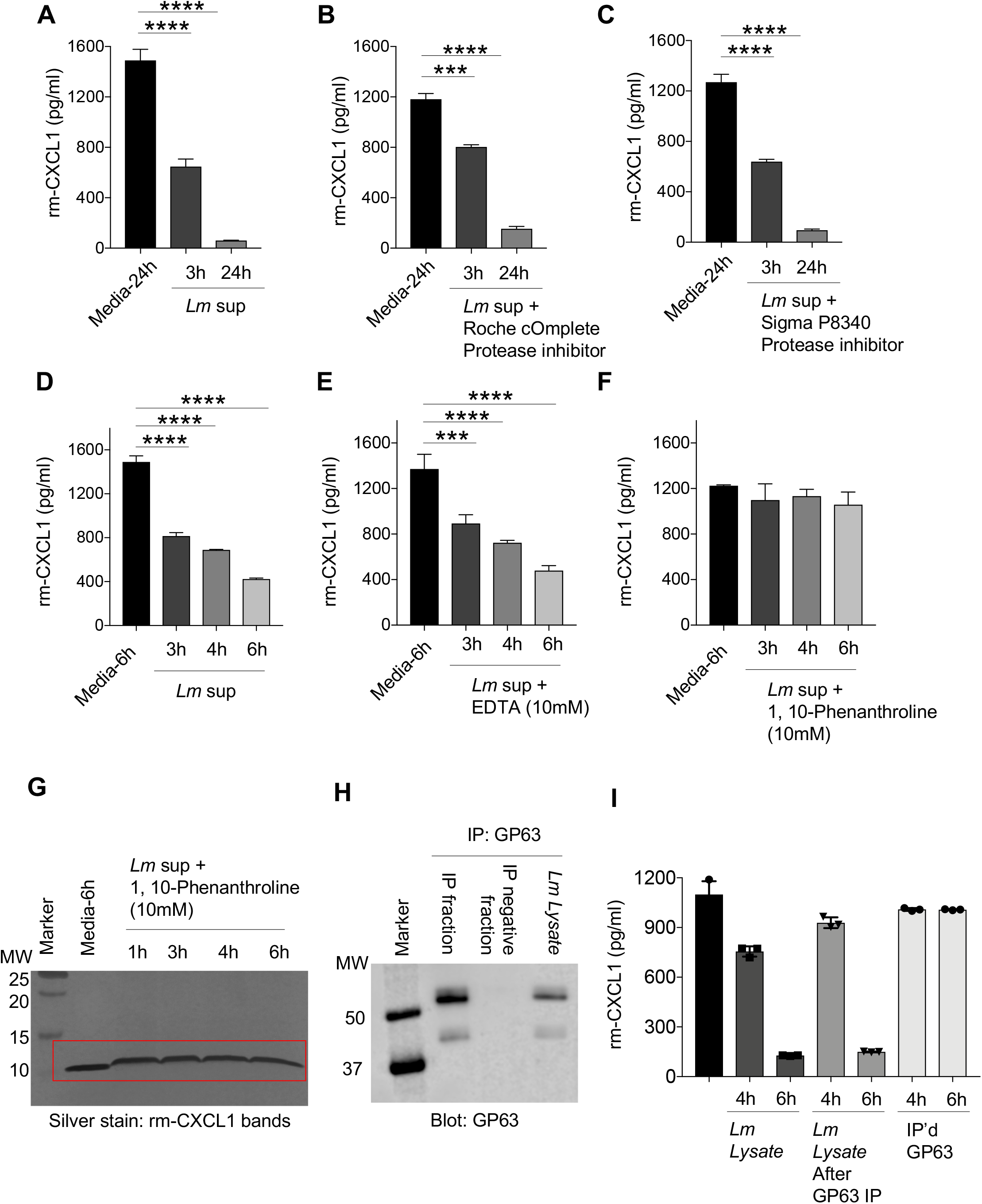
*L. major* metalloprotease cleaves CXCL1. (A) Recombinant CXCL1 was incubated with or without *Lm* sup containing (B) Roche cOmplete protease inhibitor, or (C) Sigma P8340 protease inhibitor for indicated time-points and subjected to CXCL1 quantification using ELISA. (D) Recombinant CXCL1 was treated with *Lm* sup in th presence of (E) EDTA or (F) 1,10-Phenanthroline for indicated time-points and CXCL1 degradation determined by ELISA. (G) Recombinant CXCL1 was treated with *Lm* sup in the presence of 1,10-Phenanthroline for indicated time-points and CXCL1 degradation determined by silver staining. (H) *Lm* lysate was immunoprecipitated with GP63 antibody and Western blot was carried out to evaluate the levels of GP63 protein. (I) The fractions from (H) were incubated with rm-CXCL1 for indicated time-points and ELISA was performed to quantify the levels of CXCL1. Data are representative of at least three independent experiments. Results are represented as mean ± SEM. \*\*\**P*<0.001, \*\*\*\**P*<0.0001.

Several studies have shown that 1,10-Phenanthroline inhibits GP63, a Zn^2+^ metalloprotease present on all *Leishmania spp* and is often used in biochemical assays to inhibit non-specific proteolytic activity of GP63 [51]. To determine whether GP63 was involved in specific cleavage of rm-CXCL1, we immunoprecipitated (IP) GP63 from *Lm* lysate using anti-GP63 antibody (Fig. 6H). Immunoblotting the GP63+ve and GP63-ve IP fractions for GP63 showed that GP63 was exclusively present in the IP+ve fraction and not present in the IP-ve fraction demonstrating successful immunoprecipitation of GP63 (Fig. 6H). More importantly, when these fractions were cultured with rm-CXCL1, the IP-ve (*Lm* lysate lacking GP63) but not the IP+ve (i.e. GP63) fraction degraded rm-CXCL1 suggesting a role for GP63-independent metalloprotease in the specific degradation of rm-CXCL1 (Fig. 6I).

### Synthetic peptide spanning the CXCL1 cleavage site inhibits *L. major* proteolytic activity against rm-CXCL1

Our data suggest that *L. major* Zn^2+^ metalloprotease specifically cleaves rm-CXCL1 after the K65 residue leaving a 7-mer amino acid sequence (MLKGVPK). To further examine the specificity of this *L. major* metalloprotease, we designed a blocking peptide that covered this cleavage site (i.e. the last 15 amino acid sequences of rm-CXCL1; Fig. 7A). The addition of blocking peptide was able to rescue *Lm* lysate-mediated degradation of rm-CXCL1 in a dose-dependent manner (Fig. 7B). The peptide sequence from the signal peptide region of rm-CXCL1 (peptide #1 and peptide #2, Fig. 7A) did not inhibit *Lm* lysate-mediated degradation of rm-CXCL1, demonstrating the specificity of the blocking peptide (Fig. 7B). These data demonstrate that a synthetic peptide spanning the murine CXCL1 cleavage site can competitively inhibit *L. major*-mediated rm-CXCL1 degradation.

**Figure 7:**
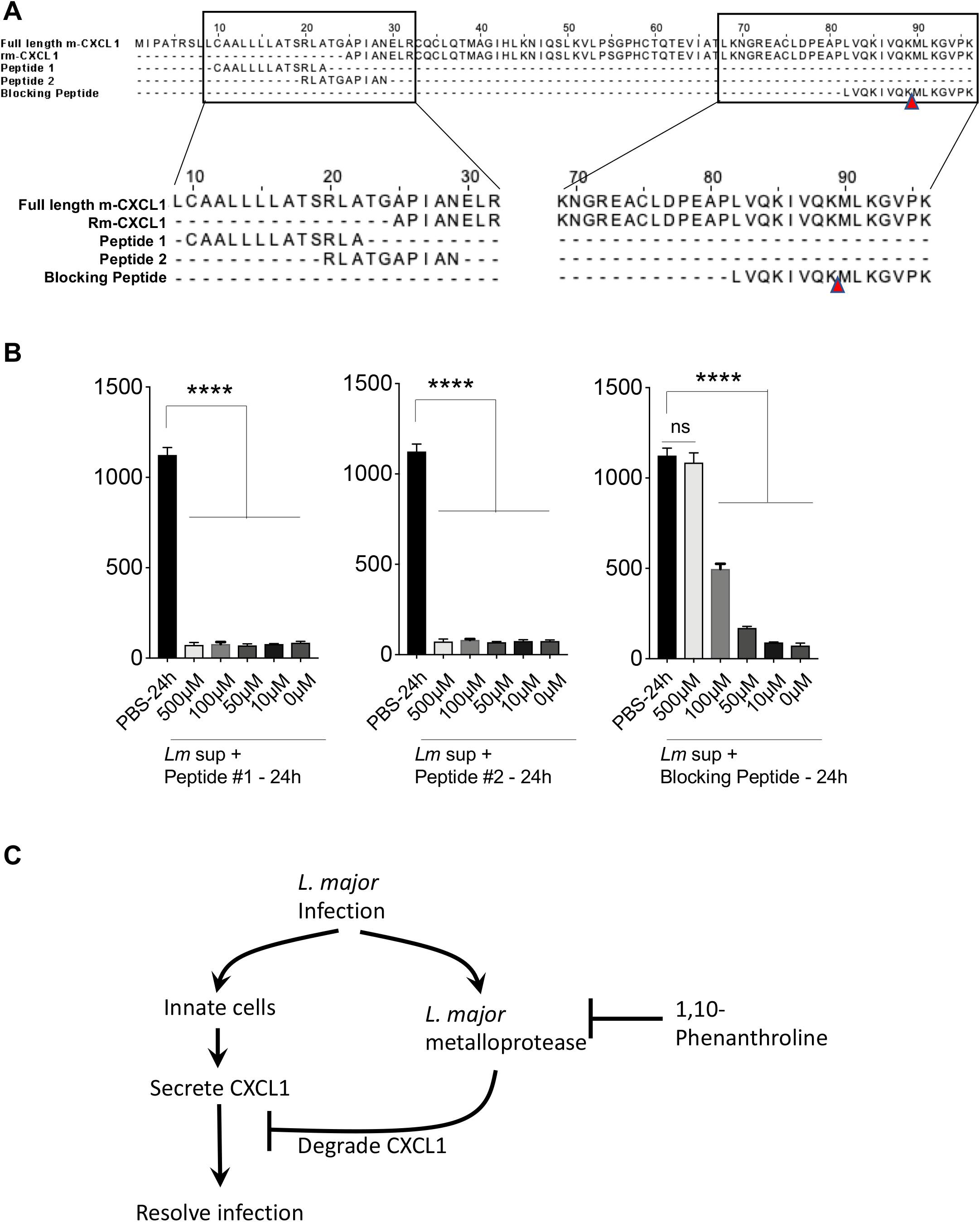
Synthetic peptide spanning rm-CXCL1 cleavage site inhibits *L. major*-mediated CXCL1 degradation. (A) Schematic of synthetic peptide generation. Alignment of full length murine CXCL1 with mature murine CXCL1 demonstrating the schematic for generation of peptide#1, peptide#2 and blocking peptide. Peptide#1 and peptide#2 were generated from the N-terminal region of full length murine CXCL1. Blocking peptide spanning the *L. major* cleavage site on murine CXCL1 were generated. *Lm* lysate-mediated degradation of rm-CXCL1 in the presence of (B) Activity of synthetic peptides from (A) in inhibiting L. major-mediated degradation of rm-CXCL1. (C) Model of *L. major* metalloprotease function on CXCL1 during infection. Data are representative of at least three independent experiments. Results are represented as mean ± SEM. \*\*\*\**P*<0.0001.

## Discussion

CXCL1 is a potent neutrophil chemoattractant and is rapidly upregulated following *Leishmania spp*. infection in the skin [36]; however, the role of CXCL1 during *Leishmania spp*. infection of mice in vivo has been understudied. Charmoy et al showed that mice deficient in CXCL1 have a slight increase in lesion size and reduced numbers of neutrophil infiltrates in chronic lesions, but the overall pathology remained similar between WT and *Cxcl1^−/−^* mice [33]. However, it should be noted that the study used the *L. major* Seidman strain (LmSd) that causes a non-healing infection in C57BL/6 mice [33]. Thus, more thorough analyses are needed to determine the precise role of CXCL1 during *L. major* infection in mice and humans.

Importantly, even depletion of neutrophils using anti-Gr1 neutralizing antibody does not always lead to increased parasitic burden, lesion or pathology [21]. While some studies with acute depletion of neutrophils reported worsened pathology during *L. major* infections, other studies have shown no role or even amelioration of disease pathology [5, 52–60]. Thus, mouse strain, *L. major* strain, route of infection and timing of CXCL1 release may all contribute to the fate of the infection and overall pathology. In line with this thought, two independent *L. major* strains were tested in our study, and both strains degraded CXCL1 in our experimental systems. Whether this degradation of murine CXCL1 is a specific feature of *L. major* or a more general feature of all *Leishmania spp*. will need further investigation.

Our initial observation demonstrated that *L. major* in culture with macrophages effectively inhibited LPS-induced CXCL1 production. Further examination showed that intracellular *L. major* within the macrophages did not inhibit LPS-induced CXCL1 production. These experiments suggest that *L. major* does not interfere with signaling pathways that promote CXCL1 production but directly regulates secreted CXCL1 in the extracellular milieu. *L. major* are present as metacyclic promastigotes in sandflies [61]. When infected into the mammalian host, *L. major* infect macrophages and resides within the vacuole where they transform into amastigotes, multiply and establish infection [62]. Because *L. major*-infected macrophages which contain amastigotes did not inhibit CXCL1 production, it could be posited that only the promastigote form of *L. major* degrades CXCL1. However, genome microarray analysis shows that the majority of genes are constitutively expressed in both *L. major* promastigotes and amastigotes (>90%) and therefore the major genes and virulent factors expressed by these different stages of *L. major* may not be different [63]. Thus, both *L. major* promastigotes and amastigotes may similarly cleave and degrade murine CXCL1 in an in vitro assay. In addition, infected macrophages can secrete exosomes containing *L. major* proteins [64, 65], which could degrade murine CXCL1. We have consistently shown that *L. major* secreted microvesicles (i.e. exosomes) contain these metalloproteases that can degrade murine CXCL1. Physiologically, *L. major* promastigotes released during a sandfly bite may act locally to limit acute CXCL1 produced in response to the infection. Once *L. major* are phagocytosed and transformed into amastigotes, they are physically separated from the CXCL1 due to their location (amastigotes are in the vacuole while CXCL1 are secreted by the macrophages and are extracellular) and may only impact CXCL1 production through the *L. major* component laden exosomes.

Our studies suggest that a yet-unknown metalloprotease from *L. major* cleaves murine CXCL1. Given that 1,10-Phenanthroline (Zn^2+^ chelator) but not EDTA (Ca^2+^ and Fe^3+^ metal ion chelator) or Marimastat (Matrix metalloprotease inhibitor) rescued *L. major*-mediated CXCL1 degradation, it is possible that the unknown metalloprotease is a Zn^2+^ metalloprotease. Our studies also demonstrate that the *L. major* metalloprotease is highly specific for murine CXCL1 in that its closest murine homolog, CXCL2, or human homologs, CXCL1, CXCL2 or CXCL8, were not degraded. We hypothesize that this particular *L. major* metalloprotease may have evolved to specifically evade host immune response in rodents. Given that human CXCL1 homologs are not susceptible to this metalloprotease, one can argue that our results may not have any importance from a public health standpoint, but we reason otherwise. Our results are highly relevant to public health because rodents are ubiquitous, serve as a reservoir for *Leishmania spp*. and we highlight a rodent-specific *L. major* evolution in targeting murine CXCL1 [66].

Our studies demonstrate that the murine CXCL1 is first cleaved at the C-terminal end after K65 releasing a 7aa residue. CXCL1 cleavage occurs as early as 30 minutes of incubation with *L. major* and by 4 hours the cleavage is complete in that only the cleaved bands are observed. Interestingly, we did not observe any accumulation of the cleaved CXCL1 band, suggesting continuous degradation of the cleaved form. While our data clearly demonstrate that the unknown *L. major* metalloprotease mediates cleavage of murine CXCL1, how the cleaved CXCL1 is further degraded will be the subject of future investigation. The cleaved murine CXCL1 (lacking the C-terminal 7 aa) may be highly unstable and undergo spontaneous degradation overtime. Alternatively, the cleaved murine CXCL1 may be susceptible to additional *L. major* proteases which promote its subsequent degradation.

In conclusion, we have identified an immune evasion strategy utilized by *L. major* that is highly specific to murine CXCL1 (Fig. 7C). Specifically, *L. major*-associated metalloprotease cleaved murine CXCL1 at K65 residue and released a C-terminal 7 amino acid fragment to promote its degradation. Finally, we have designed a peptide spanning the cleavage site of CXCL1 that inhibited murine CXCL1 cleavage by *L. major*. Our study altogether uncovered an immune evasion strategy employed by *L. major* that may have evolved in rodents and highlights how parasites may utilize diverse immune evasion strategy to establish infection within its diverse mammalian hosts.

## Methods

### Ethics statement

Experimental procedures that utilized mice were all approved by the University of Iowa Animal Care and Use Committee (Approved Animal Protocol # 7042004 – PI Dr. Prajwal Gurung) and performed in accordance to the Office of Laboratory Animal Welfare guidelines and the PHS Policy on Humane Care and Use of Laboratory Animals.

### BMDM culture

BMDMs were prepared as described previously (32). Briefly, bone marrow cells were harvested from the hind limbs of BALB/c mice (Jackson Laboratory, Stock No. 000651) and cultured in L cell-conditioned IMDM medium supplemented with 10% FBS, 1% nonessential amino acid, and 1% penicillin-streptomycin for 5-7 d to differentiate into macrophages. BMDMs were counted and seeded at 1 × 10^6^ cells in 12-well cell culture plates in IMDM media containing 10% FBS, 1% nonessential amino acids, and 1% penicillin-streptomycin. BMDMs were primed with LPS (20 ng/ml) and infected with 20 MOI *L. major* promastigotes for 24 and 48 hours (Supplemental Figure 1). For some experiments, BMDMs were LPS primed to generate supernatant containing cytokines for in vitro biochemical analysis with *L. major* supernatant (*Lm* sup) and *L. major* lysate (*Lm* lysate). BMDMs were also treated with rm-CXCL1 (20 ng/ml) or rm-CXCL1+Lm sup preparations in some experiments.

### L. major culture

*L. major* strains WHOM/IR/-173 [67] and IA0 [47] were grown in T-25 flasks with M199 media supplemented with 10% FBS, 5% HEPES and 1% penicillin-streptomycin at room temperature. BMDMs were infected with 20 MOI of *L. major* promastigotes for 48 hours. Conditioned *L. major* supernatant (*Lm* sup) was prepared after *L. major* reached the stable growth phase at approximately 20 × 10^6^ / ml. Following centrifugation (3000×g) to pellet *L. major*, supernatants were collected and filtered using a 0.2μm vacuum filter to harvest *Lm* sup. *Lm* lysate was prepared by collecting the *L. major* pellet, washing 3 times with PBS followed by 3 freeze-thaw cycles of 7 × 10^9^ *L. major* /ml in PBS. For the preparation of *L. major* exosomes, promastigotes were grown to stable phase and changed to serum-free media and incubated at room temperature overnight. Supernatant was obtained as above and then separated using a 100kDa molecular weight Amicon ultra-15 centrifugal filter (MilliporeSigma, Burlington, MA). The concentrated vesicles were washed twice with PBS by ultracentrifugation at 110,000xg at 4°C for 1 hour (Beckman Optma MAX-XP Ultracentrifuge, rotor TLA 120.2; Beckman Coulter Inc., USA). The final pellet (exosomes) was resuspended in PBS and used in in vitro biochemistry assays.

### Electrophoresis and in vitro biochemistry

For in vitro reactions, 125ng of total recombinant protein was separated on 4-20% or 10-20% Novex WedgeWell Tris-Glycine precast gels (Thermofisher, Waltham, MA). For silver stain, the Pierce Mass Spec compatible silver stain kit was used per manufacturer’s instructions (Thermofisher). Synthetic peptides were generated and purchased from Pierce Biotechnology (Thermofisher) and are composed of the following sequences: peptide#1: CAALLLLATSRLA; peptide#2: RLATGAPIAN; blocking peptide: LVQKIVQKMLKGVPK. For Western blot analysis a semi-dry transfer was completed using trans-blot turbo system (Bio-Rad, Hercules, CA) with Immobilon PVDF membrane. The membranes were blocked in 5% non-fat milk for 1 hour at room temperature. Membranes were probed with sheep polyclonal antibody Sp180 raised against *L. chagasi* promastigote GP63 at a concentration 1:1500 incubated overnight at 4°C. The membranes were then probed with HRP-tagged secondary antibodies at room temperature for 1 hour and developed using Immobilon Forte Western HRP Substrate (MilliporeSigma) and imaged with an Odyssey Fc Infrared Imaging System (LI-COR Bioscience, Lincoln, NE). Immunoprecipitation was performed using Protein G Dynabeads and Leishmania GP63 monoclonal antibody clone 96-126 (Thermofisher). As instructed by the manufacturer, 50μL beads were washed in PBS with 0.02% Tween 20 and bound to 5μg antibody for at least 10 minutes. Following a wash, the antibody bound beads were incubated with *Lm* lysate for another 10 minutes and finally either antigen-antibody binding was negated by acidic (pH 2.8) 50mM Glycine treatment or bead bound antibody-antigen complexes were used directly for biochemistry.

### Mass spectrometry

#### In-gel trypsin digestion

The gel was stained using a Pierce mass spec compatible silver stain kit (Thermofisher) per manufacturer directions. A procedure slightly modified than that described by Yu et al [68] was used for in-gel digestion. Briefly, the targeted protein bands from SDS-PAGE gel were manually excised, cut into 1mm^3^ pieces, and washed in 100mM ammonium bicarbonate:acetonitrile (1:1, v/v) and 25mM ammonium bicarbonate /acetonitrile (1:1, v/v), respectively, to achieve complete de-staining. The gel pieces were further treated with acetonitrile (ACN), to effectively “dry” the gel segments and then reduced in 50μl of 10mM DTT at 56 °C for 60 min. Gel-trapped proteins were alkylated with 55mM chloroacetamide (CAM) for 30 min at room temperature. The gel pieces were washed with 25mM ammonium bicarbonate: acetonitrile (1:1, v/v) twice to remove excess DTT and CAM. 50μl of cold trypsin solution at 10ng/μl in 25mM ammonium bicarbonate was then added to the gel pieces and they were allowed to swell on ice for 60 min. Digestion was conducted at 37°C for 16 h. Peptide extraction was performed three times, adding 100μL of 50% acetonitrile/0.1% formic acid for 0.5 h, combining the supernatants. The combined extracts were concentrated in a lyophilizer and rehydrated with 15 μl of mobile phase A.

#### LC-MS/MS

Mass spectrometry data were collected using an Orbitrap Fusion Lumos mass spectrometer (Thermofisher) coupled to an Easy-nLC-1200™ System (Proxeon P/N LC1400). The autosampler is set to aspirate 3μl (estimated 0.3ug) of reconstituted digest and load the solution on a 2.5cm C18 trap (New Objective, P/N IT100-25H002) coupled to waste, HV or analytical column through a microcross assembly (IDEX, P/N UH-752). Peptides are desalted on the trap using 16μl mobile phase A for 4 min. The waste valve is then blocked and a gradient begins flowing at 0.4μl/min through a self-packed analytical column, 10cm in length × 75μm id. The fused silica column was tapered from 100μm ID (Polymicro) to ~8μm at the tip using a Sutter P-2000 laser puller then packed with 2.7micron Halo C18 particles using a He-pressurized SS cylinder. Peptides were separated in-line with the mass spectrometer using a 70 min gradient composed of linear and static segments wherein buffer A is 0.1% formic acid and B is 95%ACN, 0.1% formic acid. The gradient begins first hold at 4% for 3 min then makes the following transitions (%B, min): (2, 0), (35, 46), (60, 56), (98, 62), (98, 70).

#### Tandem mass spectrometry on the Thermo Q-Exactive hf

Data acquisitions begin with a survey scan (m/z 380 −1800) acquired on a Q-Exactive Orbitrap mass spectrometer (Thermofisher) at a resolution of 120,000 in the off-axis Orbitrap segment (MS1) with automatic gain control (AGC) set to 3E06 and a maximum injection time of 50 ms. MS1 scans were acquired every 3 sec during the 70-min gradient described above. The most abundant precursors were selected among 2-6 charge state ions at a 1E05 AGC and 70 ms maximum injection time. Ions were isolated with a 1.6 Th window using the multi-segment quadrupole and subjected to dynamic exclusion for 30 sec if they were targeted twice in the prior 30 sec. Selected ions were then subjected to high energy collision-induced dissociation (HCD) in the ion routing multipole (IRM). Targeted precursors were fragmented by (HCD) at 30% collision energy in the IRM. HCD fragment ions were analyzed using the Orbitrap (AGC 1.2E05, maximum injection time 110 ms, and resolution set to 30,000 at 400 Th). Both MS2 data were recorded as centroid and the MS1 survey scans were recorded in profile mode.

#### Proteomic searches

Initial spectral searches were performed with both Mascot version 2.6.2 (MatrixScience) and Byonic search engines (Protein Metrics ver. 2.8.2). Search databases were composed of the Uniprot KB for species 10090 (mouse) downloaded February 6, 2016 containing 58436 sequences. In either search, an equal number of decoy entries were created and searched simultaneously by reversing the original entries in the target database. Precursor mass tolerance was set to 5 ppm and fragments were searched at 10 ppm. A fixed 57 Da modification was assumed for cysteine residues while variable oxidation was allowed at methionine. A variable GG modification at lysine was set to monitor ubiquitylation and potential phosphorylation was accessed at Ser and Thr residues. The false discovery rate was maintained at 1% by tracking matches to the decoy database. Both Mascot and Byonic search results were combined and validated using Scaffold ver. 4.8.5 (Proteome Software). Protein assignments required a minimum of two peptides established at 70% probability (local FDR algorithm) and an overall 95% protein probability (assigned by Protein Prophet). Approximately 300 protein families (including common contaminants) were assigned at a total FDR to 1.2%. Proteins were annotated with GO terms from goa_uniprot_all.gaf downloaded on May 3, 2017.

### Immunoassays and recombinant proteins

Cytokine ELISAs and multiplex immunoassays were performed according to manufacturer instructions. Mouse and human CXCL1 and CXCL2 as well as human IL-8 (CXCL8) were obtained from R&D Systems (Minneapolis, MN) and multiplex CXCL1, IL-6, and TNF were completed using ProcartaPlex assays (Thermofisher) which were run on the BioRad Bio-Plex (Luminex, Austin, TX). We obtained our recombinant proteins from Tonbo biosciences which included mouse CXCL1, CXCL2, and TNF as well as human CXCL1, CXCL2, and CXCL8.

### PCR methods

RNA was extracted from BMDM cultures in TRIzol reagent (Thermofisher) followed by chloroform extraction and isopropanol precipitation. The extracted RNA was reverse-transcribed into cDNA by using qScript Supermix (Quanta Biosciences, Beverly, MA). Real-time quantitative PCR was performed on Eppendorf realplex EPgradient S Mastercycler (Eppendorf, Germany) using PerfeCTa SYBR Green SuperMix ROX (Quanta Biosciences) and the appropriate primers. The sequences of the quantitative RT-PCR primers are as follows: *mCXCL1*: 5’-CAATGAGCTGCGCTGTCAGTG-3’, 5’-CTTGGGGACACCTTTTAGCATC-3’; *mTnf*: 5’-CATCTTCTCAAAATTCGAGTGACAA-3’, 5’-TGGGAGTAGACAAGGTACAACCC-3’; *mIL-6*: 5’-CAAGAAAGACAAAGCCAGAGTC-3’, 5’-GAAATTGGGGTAGGAAGGAC-3’. Threshold cycle Ct values were normalized to GAPDH, and gene fold change was determined by the relative comparison method, relative to the 0 h time point.

### Statistical analysis and alignment

GraphPad Prism 8.0 software was used for data analysis and figure presentation. Data are shown as mean ± SEM. Statistical significance was determined by *t* tests (two-tailed and Mann-Whitney) for two groups, and one-way ANOVA (with Dunnett’s or Tukey’s multiple comparisons tests) for three or more groups. For the purpose of alignment presentation, sequences were obtained from the protein manufacturer or the NCBI protein database and figures were prepared using Jalview 2 [69].

## Authorships

M.S.Y, B.P., L.M. performed the experiments. M.S.Y, M.E.W., and P.G. analyzed the data. R.M.P. performed the mass spectrometry and analysis of CXCL1 cleavage bands. M.S.Y. and P.G. wrote the manuscript with input from all authors. P.G. designed and conceptualized the study.

## Acknowledgements

We thank the University of Iowa Proteomics Core for the comparative analysis performed in the University of Iowa Proteomic facility, Directed by RMP and supported by an endowment from the Carver Foundation and by Thermo Lumos provided through an HHMI grant to Dr. Kevin Campbell. We thank Kristina Greiner for her help with scientific editing of the manuscript. We are also thankful to the members of the Inflammation Program for critical insights provided during the development of the project. This work was supported by NIAID K22AI127836 Grant, NIEHS P30ES005605 Pilot Grant, and the University of Iowa Startup funds to Dr. Prajwal Gurung.

---

**Supplemental Figure 1:**
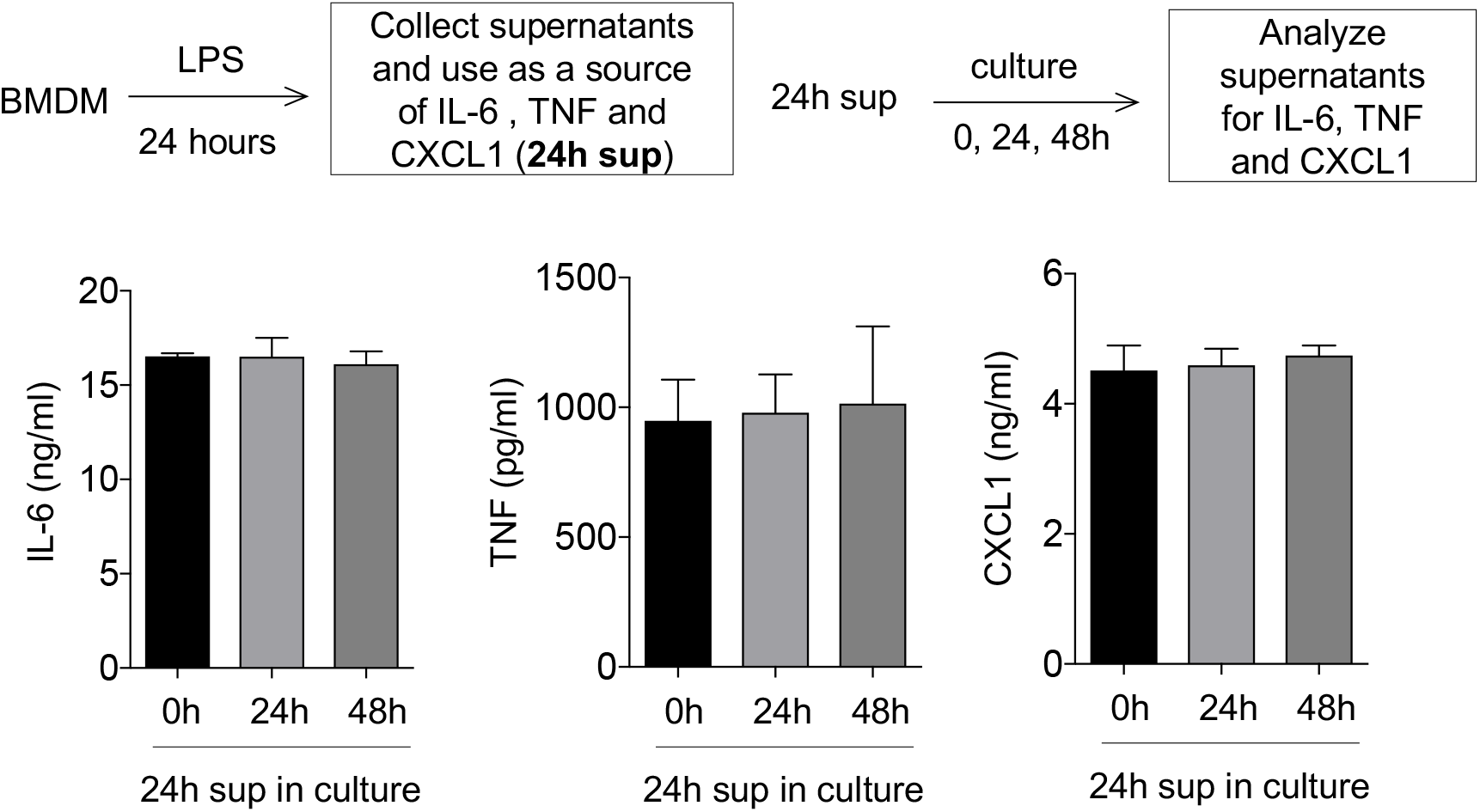
CXCL1, IL-6, and TNF secreted by LPS-activated BMDMs are stable in cell culture conditions. Conditioned supernatants from 24h LPS (20 ng/ml) stimulated BMDMs were collected and further cultured for 0, 24, or 48h and stability of CXCL1, IL-6 and TNF in the culture were determined by ELISA. Data are representative of at least three independent experiments. Results are represented as mean ± SEM.

**Supplemental Figure 2:**
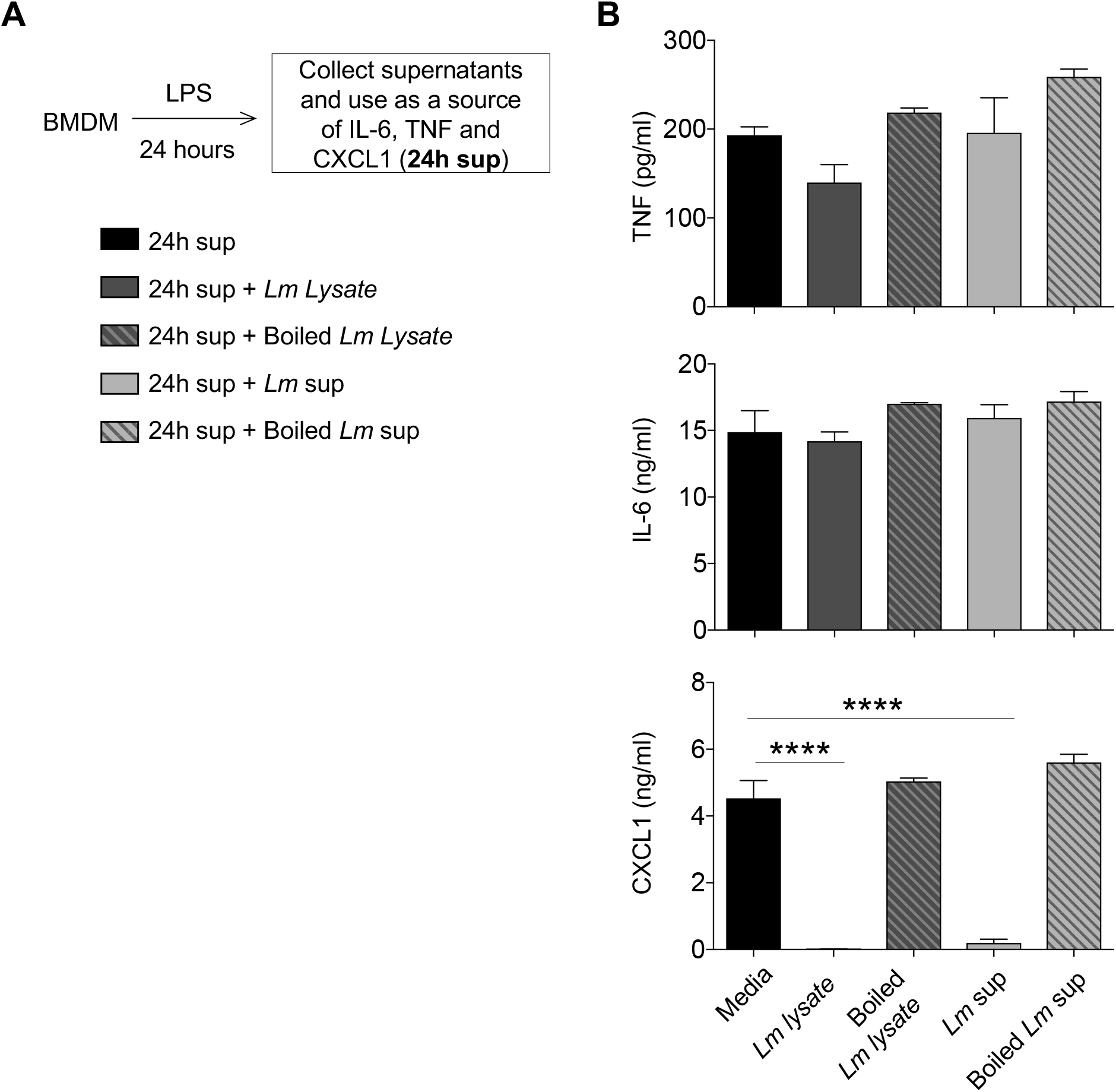
Heat inactivation of *Lm* lysate or *Lm* sup rescues CXCL1 detection. (A) Experimental Design. (B) Conditioned supernatants from 24h LPS (20 ng/ml) stimulated BMDMs were collected and subjected to 24h treatment with control or boiled (20 min at 100°C) *Lm* lysate or *Lm* sup. Levels of CXCL1, IL-6 and TNF were determined by ELISA. Results are represented as mean ± SEM. \*\*\*\**P*<0.0001.

**Supplemental Figure 3:**
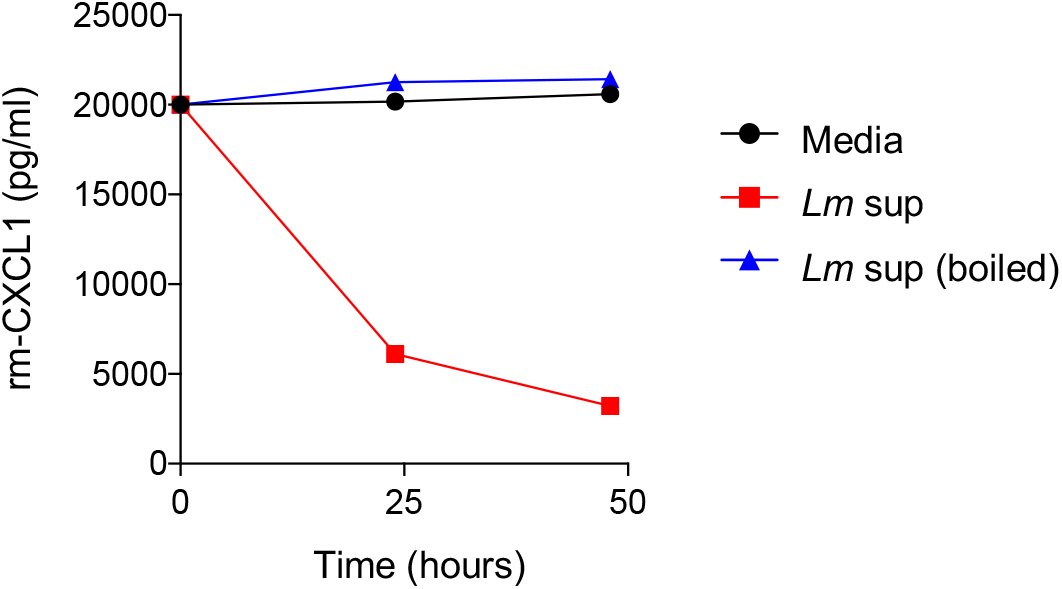
*L. major* reduces rm-CXCL1 levels in culture. Rm-CXCL1 were left alone or treated with *Lm* sup (control or boiled) for up to 48 hours in culture. The quantity of rm-CXCL1 in the culture at 0, 24 and 48 h were determined by ELISA. Data are representative of at least three independent experiments. Results are represented as mean ± SEM.

**Supplemental Figure 4:**
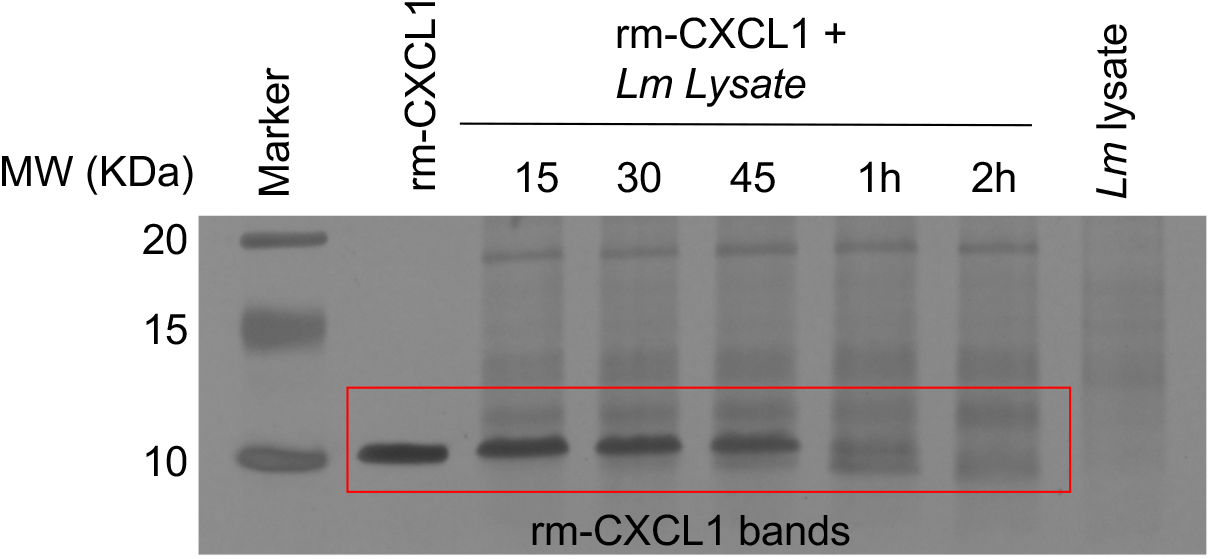
Acute time course of *L. major*-induced CXCL1 cleavage. Rm-CXCL1 were left alone or treated with *Lm* lysate for acute time points as indicated to determine the precise time course of rm-CXCL1 cleavage. Silver staining was used to visualize rm-CXCL1. Data are representative of at least three independent experiments.

**Supplemental Figure 5:**
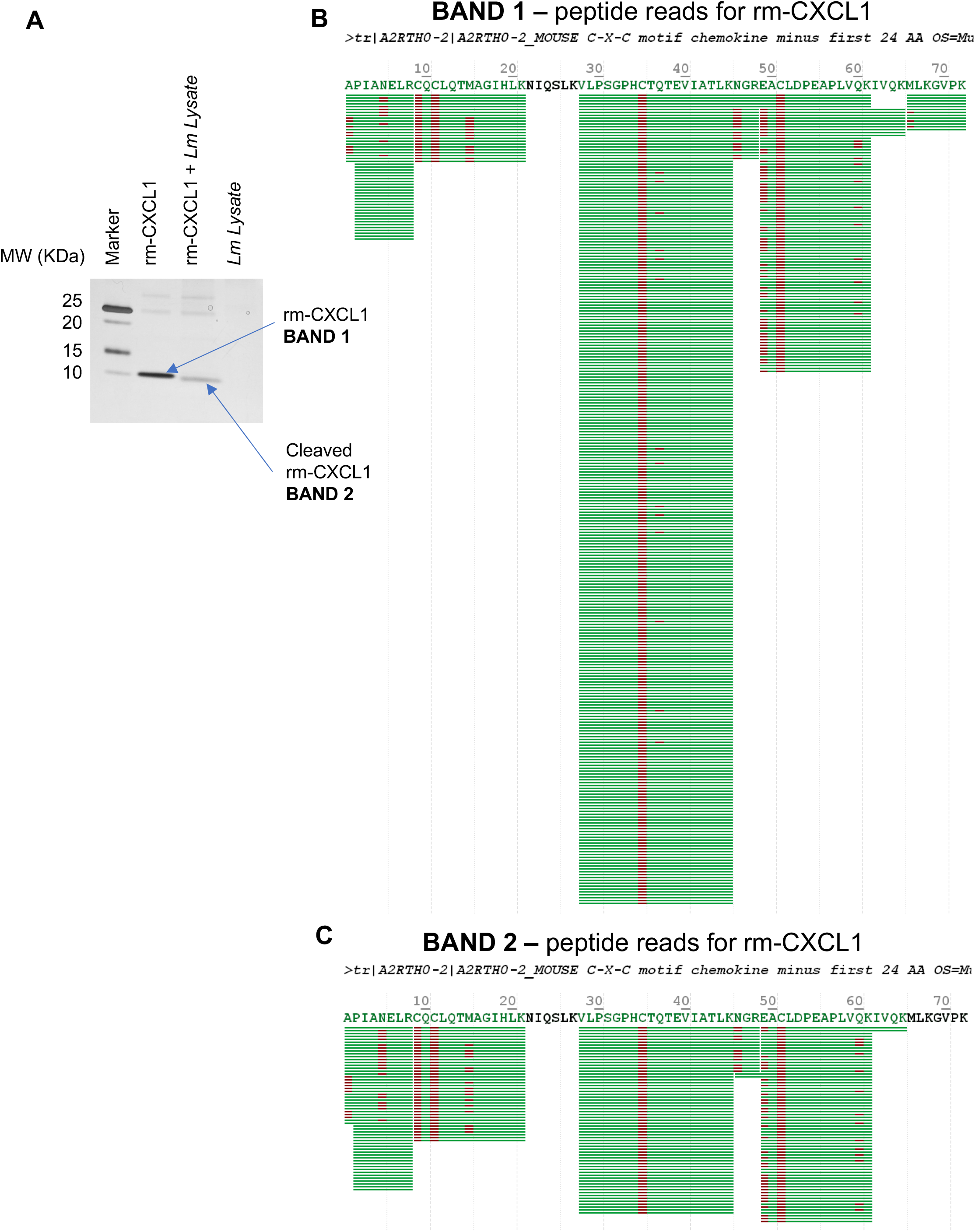
Identification of CXCL1 cleavage site and cleaved products. (A) A silver-stained SDS-PAGE showing cleavage of full length (BAND 1) and cleaved rm-CXCL1 (BAND 2). (B) Mass spectrometry analysis of BAND 1 (full length CXCL1) and tryptic peptide coverage analysis confirm a full length rm-CXCL1. (C) Mass spectrometry analysis of BAND 2 (cleaved CXCL1) and tryptic peptide coverage analysis confirm a C-terminal end cleavage after K65 residue.

**Supplemental Figure 6:**
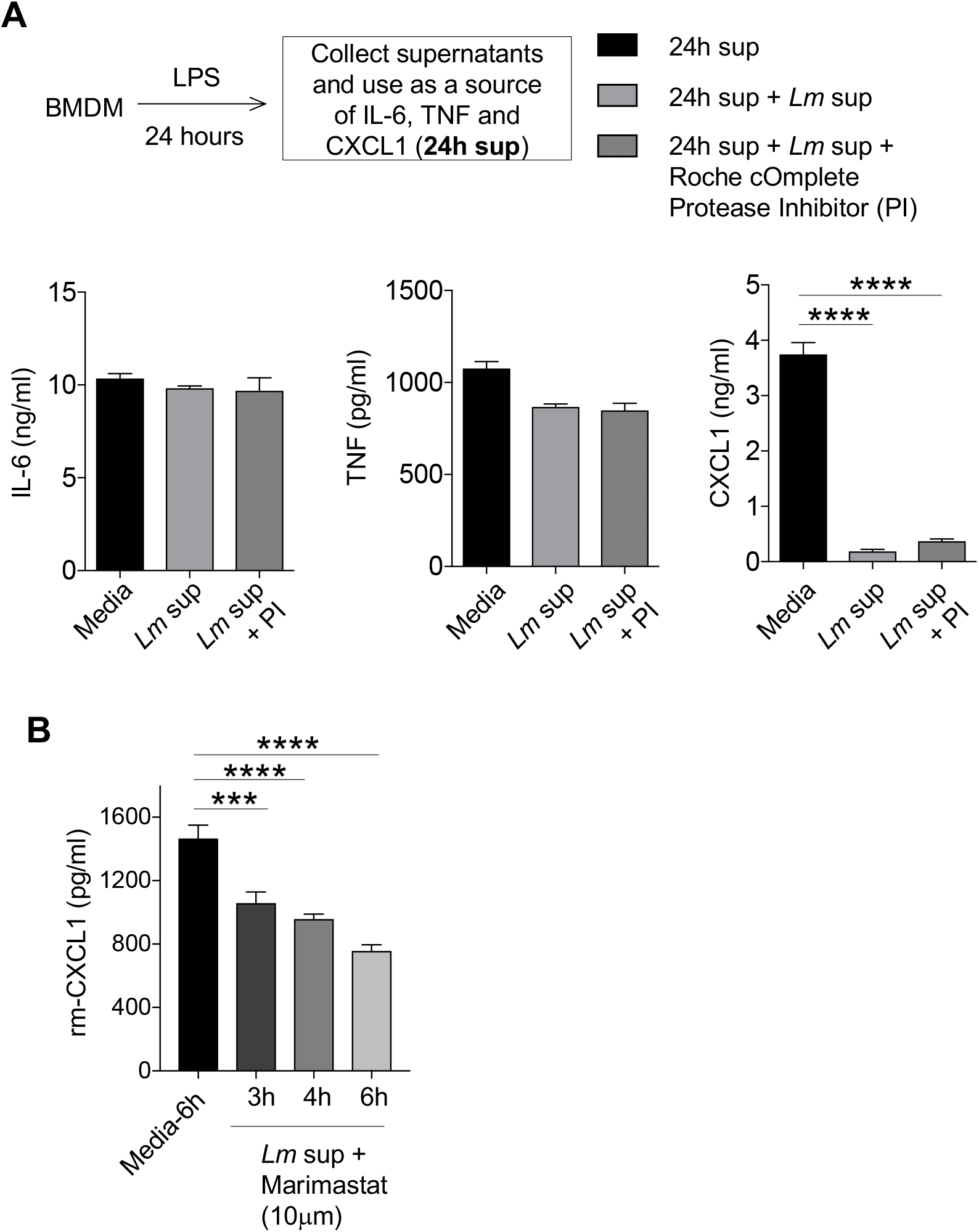
Proteolytic activity of *L. major* on CXCL1 is not prevented by Roche cOmplete Inhibitor or Marimastat. (A) Conditioned supernatants from 24h LPS (20 ng/ml)-stimulated BMDMs were collected and treated with *Lm* sup in the presence or absence of Roche cOmplete protease inhibitor. Levels of CXCL1, IL-6 and TNF were determined by ELISA. (B) Rm-CXCL1 was treated with *Lm* sup in the presence or absence of Marimastat (10μM) for indicated time-points and ELISA was performed to evaluate the levels of CXCL1. Data are representative of at least three independent experiments. Results are represented as mean ± SEM. \*\*\**P*<0.001, \*\*\*\**P*<0.0001.

